# Rainbowfish: A Succinct Colored de Bruijn Graph Representation

**DOI:** 10.1101/138016

**Authors:** Fatemeh Almodaresi, Prashant Pandey, Rob Patro

## Abstract

The colored de Bruijn graph— a variant of the de Bruijn graph which associates each edge (i.e., k-mer) with some set of colors — is an increasingly important combinatorial structure in computational biology. Iqbal et al. demonstrated the utility of this structure for representing and assembling a collection (population) of genomes, and showed how it can be used to accurately detect genetic variants. Muggli et al. introduced VARI, a representation of the colored de Bruijn graph that adopts the BOSS representation for the de Bruijn graph topology and achieves considerable savings in space over `Cortex`, albeit with some sacrifice in speed. The memory-efficient representation of VARI allows the colored de Bruijn graph to be constructed and analyzed for large datasets, beyond what is possible with `Cortex`.

In this paper, we introduce Rainbowfish, a succinct representation of the color information of the colored de Bruijn graph that reduces the space usage even further. Our representation also uses BOSS to represent the de Bruijn graph, but decomposes the color sets based on an equivalence relation and exploits the inherent skewness in the distribution of these color sets. The Rainbowfish representation is compressed based on the 0th-order entropy of the color sets, which can lead to a significant reduction in the space required to store the relevant information for each edge. In practice, Rainbowfish achieves up to a 20 × improvement in space over VARI. Rainbowfish is written in C++11 and is available at https://github.com/COMBINE-lab/rainbowfish.

## 1 Introduction and Related Work

This paper proposes a new representation of the colored de Bruijn graph. The colored de Bruijn graph is a variant of the de Bruijn graph where each edge (i.e., *k*-mer) is associated with some set of colors. Here, each color is used to encode the source of the corresponding *k*-mers (e.g., different source genomes, transcriptomes, sequenced samples, etc.). From this perspective, it is a flexible and powerful combinatorial structure for representing a collection of sequences while maintaining the identity of each. This structure gained popularity in the work of Iqbal et al. [6], which demonstrated the utility of the colored de Bruijn graph for representing and assembling a collection (population) of genomes, and for detecting both simple and complex genetic variants with high accuracy. Analysis of the colored de Bruijn graph exhibits particular promise for analyzing complex population-level variation, since topological structures (e.g., bubbles) can be associated with variation in the underlying sub-populations. The representation adopted by Iqbal, as implemented in the tool `Cortex`, is optimized for speed, and so requires a considerable amount of memory to represent both the topology of the de Bruijn graph and the colors associated with each edge.

The memory usage of the colored de Bruijn graph representation adopted in `Cortex` precludes this approach from being adopted when the underlying genomes and color sets become too large. In order to overcome such limitations, Muggli et al. [10] introduced the VARI representation of the colored de Bruijn graph. This approach sacrifices some of the speed of the `Cortex` representation for a considerable reduction in the required space. VARI achieves this space savings in two ways. First, rather than using a hash-table-based representation of the de Bruijn graph topology, it adopts the highly-efficient BOSS representation. The BOSS [1] representation (named based on the initials of the authors) makes use of the FM index to encode the topology of the de Bruijn graph. BOSS uses 4*N* + *o(N)* bits to represent a de Bruijn graph with *N* edges (empirically, this often works out to be as few as 4-6 bits per edge).

VARI couples the BOSS representation of the de Bruijn graph topology with a compressed representation of the color information. By it’s nature, BOSS assigns to every de Bruijn graph edge a distinct rank in the range [0, *N*). So, VARI represents the color information as a *N × C* bit matrix where *C* is the number of input colors. Conceptually, each of the *N* rows of this matrix is simply a bit vector that encodes which of the *C* colors label the corresponding edge. To reduce the space required to store this color information, VARI concatenates these rows into a single (*N × C*) × 1 vector and stores them in an Elias-Fano encoded bit vector, allowing for a (sometimes substantial) reduction in the size while still enabling efficient point queries (i.e., is a particular edge labeled with a given color?). Muggli et al. [10] demonstrate that the VARI representation can be built on data sets consisting of large numbers of *k*-mers, large input color sets, or both. Specifically, the space efficiency of VARI makes it possible to build and query the colored de Bruijn graph on datasets that are orders of magnitude larger than what is possible with `Cortex`. This is an exciting development that opens up this methodology for increasingly large-scale analysis.

Though VARI achieves a substantial improvement in space over `Cortex`, there is still a considerable amount of redundancy present in its representation. Both of these systems represent the color set corresponding to each *k*-mer independently of other *k*-mers. However, many of the possible subsets of colors do not occur in practice, and there is an inherent (often extreme) skewness in the distribution of the color sets that do appear. Hence a considerable amount of redundant information can be present when the color set for each *k*-mer is represented independently. It becomes even more important to exploit this skewness for large metagenomic datasets because the space usage of VARI for these datasets can become impractical.

In this paper, we introduce a succinct representation, called Rainbowfish, of the color sets associated to each edge in the de Bruijn graph. We also adopt the BOSS representation of the de Bruijn graph topology, and focus, specifically, on how to concisely represent the color information. Rainbowfish’s colored de Bruijn graph representation is entropy compressed and exploits the high skewness present in the distribution of color sets. By exploiting a more efficient decomposition of the set of present colors (i.e., in terms of equivalence classes), we achieve a considerable reduction over the space required by VARI (up to 20× depending on the dataset), while still retaining efficient (i.e., constant time) queries.

## 2 Background and definitions

Rainbowfish is a succinct representation of the color information, and uses rank and select operations to lookup the color class corresponding to *k*-mers in the de Bruijn graph. Here, we briefly recapitulate the definition of a succinct data structure and the rank and select operations.

A ***succinct data structure*** consumes an amount of space that is close to the information-theoretic optimum. More precisely, if *Z* denotes the information-theoretic optimal space usage for a given data structure, then a succinct data structure uses *Z + o(Z)* space [8].

***rank*** and ***select*** [7] are operations that are commonly used for navigating within succinct data structures. For a bit vector *B[0,…, n— 1]*, RANK(*j*) returns the number of 1s in the prefix *B[0,…,j]* of B. SELECT(*r*) returns the position of the *r*th 1, that is, the smallest index *j* such that RANK(*j*) = *r*. For example, for the 12-bit vector B[0,…, 11] =100101001010, rank(5) = 3, because there are three bits set to one in the 6-bit prefix *B[0,…*, 5] of B, and select(4) = 8, because B[8] is the fourth 1 in the bit vector.

## 3 Method

In this section we first describe the design of Rainbowfish. We then analyze the space usage and provide a lower bound for the representation of sets of colors given a ranking of de Bruijn graph edges. Finally, we discuss the Rainbowfish implementation.

### 3.1 Design

Rainbowfish’s compact representation of color information is based on two particular observations. First, it is often the case that many of the *k*-mers in a colored de Bruijn graph share the same set of colors. More formally, we define an equivalence relation ~ over the set of *k*-mers in the de Bruijn graph. Let `Col(.)` denote the function that maps each *k*-mer to its corresponding set of colors. We say that two *k*-mers are color-equivalent (i.e., *k_1_*~ *k_2_*) if and only if `Col`(*k*_1_) = `Col`(*k*_2_). We will refer to the set of colors shared by the *k*-mers related by ~ as a *color class*. If *C*, the number of input colors, is large, it is often the case that the number of distinct color classes is far less than the number of possible color classes (which is bounded above by min(*N, 2^C^*)).

Second, it is often the case that the frequency distribution of color classes is far from uniform. Hence, it will often be useful to record a frequently occurring color class using a short description (i.e., a small number of bits) while reserving larger descriptions for less frequent color classes.

The design of Rainbowfish is motivated by the above observations. Instead of storing the color set for each *k*-mer separately, Rainbowfish stores each distinct color class only once and assigns to each distinct class a variable-length label (which, practically, is much smaller than the unary encoding of the color class itself). It then stores, for each *k*-mer, the label of the color class to which it belongs.

The approach we use to assign variable-length labels to color classes is similar in spirit to the construction of a Huffman code, where the message is a string of color class symbols. However, we do not build a prefix code, and instead opt to store an additional bit vector to allow the efficient selection of an arbitrary label from the list. We generate the labels according to the following procedure. We first sort, in descending order, all the color classes based on their frequency (i.e., the number of *k*-mers in this color equivalence class). We then assign labels to each color class starting from the class with the largest cardinality, so that the color class represented by the most frequent label will have the shortest label length etc.

The color class representation in Rainbowfish has three components. Rainbowfish stores the mappings between labels and color classes in an ***equivalence class table (ECT)***. As labels are assigned sequentially, this is simply an array of bit vectors encoding the corresponding color sets. Apart from the equivalence class table, Rainbowfish maintains two bit vectors, a ***boundary bit vector (BBV)*** and a ***label bit vector(LBV)***.

All color classes are stored in the equivalence class table (with their corresponding labels implicitly being their position). However, we now need to store a mapping from *k*-mers to the variable-length labels. Rainbowfish stores variable-length labels corresponding to each k-mer in the label bit vector. The labels are stored in the order in which *k*-mers are stored in the de Bruijn graph representation. Specifically, the *k*-mers are stored in the rank order induced by BOSS. However, since these labels are variable-length, we can not directly read the label corresponding to the k-mer of a specific rank, since we don’t know where such a label begins or how long it is.

To address this, Rainbowfish maintains another bit vector — the boundary bit vector (BBV) — to mark the boundary of each variable-length label in LBV. The boundary bit vector is the same size as the label bit vector and has a bit set to 1 at each index where a new label starts in the LBV. Thus, the starting position for the label corresponding to the rth k-mer can be obtained by issuing a SELECT(r) query on BBV, and the length of this label can be obtained by simply scanning BBV until we encounter the next set bit. Note; since the number of distinct labels is bounded by the number of *k*-mers in the dataset, in practice, the next set bit will always occur within one machine word length of where the label begins (i.e., we never have > 2^64^ distinct labels).

Figure 1 shows how the color classes are represented in Rainbowfish. To perform a query for the color class corresponding to a k-mer in the colored de Bruijn graph, we first get the rank *r* of the k-mer in the de Bruijn graph. We then perform a select operation using *r* on the boundary bit vector. The result of the select operation *i* is the start index of the label of the color class in the label bit vector to which the k-mer belongs. To find the length of the label we determine the index *i* of the next bit set in the boundary bit vector using the TZCNT instruction. Using *i* and *i’* we retrieve the label from the label bit vector, and using the label we lookup the corresponding color class in the equivalence class table.

**Figure 1.**
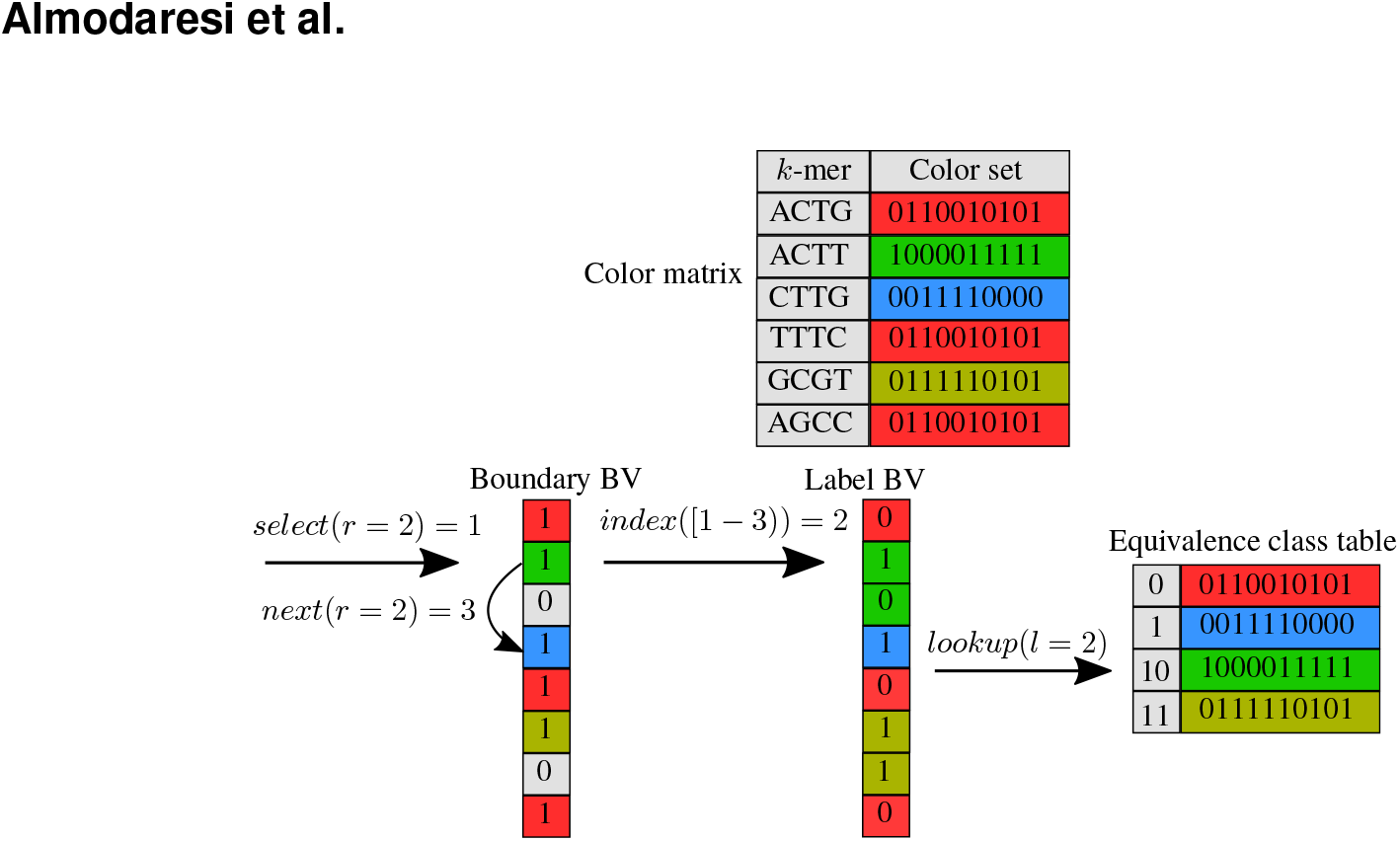
The representation of color information in Rainbowfish. The “Color Matrix” at the top represents 6 distinct 4-mers, each assigned a color set. 3 of these 4-mers (ACTG, TTTC, AGCC) have the same color class, labeled 0, and the other 3 (CTTG, ACTT, and GCGT) each have color classes labeled 1, 10, and 11 respectively. To retrieve the color set for a *k*-mer, we first perform select on the boundary bit vector (BBV) using rank *r* of the corresponding edge (*k*-mer). This returns the label’s starting position, *i*. We then look for the next set bit BBV to find the label’s ending position, *j*. Then, we fetch the label at indices *i* to *j* in label bit vector (LBV). Finally, we lookup the label *l* in the equivalence class table (ECT) and return the color class corresponding to the label.

### 3.2 Space analysis

The color class representation in Rainbowfish is entropy compressed, i.e., the space is bounded by the entropy *(H(X_c_))* of the color class distribution. For a dataset in which number of *k*-mers belonging to each distinct color class are similar, the entropy of the color class distribution will be high. On the other hand, if most of the *k*-mers in a dataset belong to a small number of distinct color classes, the entropy of the color class distribution will be low.

#### Lemma 1.

*The size of each color class label is bounded by log_2_ M bits, where M is the total number of distinct color classes. For a dataset with N distinct k-mers coming from C input samples (i.e., colors), we have that M≤min(N, 2^C^)*.

#### Theorem 2.

*Given an ordering of edges (or k-mers) in a de Bruijn graph, the space needed by Rainbowfish to represent a color class attached to each edge is O(MC + NH(X_c_)) bits, where M is the number of distinct color classes, C is the number of colors, N is the number of distinct k-mers, and 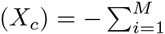 (i.e., order-0 or Shannon’s entropy) is the entropy over random variable X_c_ distributed according to the frequency distribution of the color classes*.

**Proof.** The space needed by Rainbowfish can be analyzed as follows. There are three bit vectors in Rainbowfish, the equivalence class table, label bit vector, and boundary bit vector. To store an equivalence class table containing *M* distinct color classes each having *C* colors we need *MC* bits.

To store a label bit vector (as stated in Theorem 1), for N *k*-mers, where each label corresponds to one of the *M* distinct color classes, takes *N* log_2_ *M* bits. However, as explained in Section 3.1, in Rainbowfish we assign (optimal) variable-length labels based on the frequency of color classes. Therefore, the space needed to store the label bit vector is dependent on the 0th-order entropy of the color class variable, *H(X_c_)*, and the size of the label bit vector is upper bounded by *N* log_2_ *M*. The boundary bit vector has the same number of bits as the label bit vector.

### 3.3 Lower bound for color representation

We now provide a lower bound to store a color class representation for a set of edges in a colored de Bruijn graph. In the color class representation, the equivalence class table takes *MC* bits to store *M* bit vectors each having *C* bits, which is optimal. The other two bit vectors, the boundary and label bit vector, map *k*-mers given an ordering in the de Bruijn graph to their corresponding color classes. The theorem below gives the lower bound to store such a mapping.

#### Theorem 3.

*The lower bound to represent a mapping from an ordered list of k-mers in a de Bruijn graph to a set of color classes is log_2_ (M^N-M^.M!) bits, where M is the number of distinct color classes, N is the number of edges, and for a dataset with N distinct k-mers coming from C input samples (i.e., colors), we have that M≤min(N, 2^C^)*.

**Proof.** We can analyze the lower bound using a counting argument. We count the number of ways to map a set of *M* distinct color classes to a set of *N* edges. The space required to store the color class representation should be less than or equal to the space required to store these mappings.

Edges can be mapped to color classes using a surjective (onto) function. Thus, we wish to count the total number of surjections from *M* color classes to *N* edges. Rather than counting this number exactly, we instead provide a lower bound. First, we must ensure that each of the *M* color classes maps to at least one edge — so, we select a set of *M* edges and label each with a distinct color class. There are *M*! ways to assign *M* color classes to a set of *M* edges. We will then allow the remaining *N – M* edges to be colored in any possible manner. We can assign *M* colors to *N – M* edges (the remaining number) in *M^N–M^* ways. Therefore, the total number of different mappings is bounded below by *M^N–M^. M!*. To be able to represent each such mapping, and distinguish it from the others, we need at least log_2_ *(M^N-M^ . M!)* bits.

The lower bound can be expanded using Sterling’s approximation as

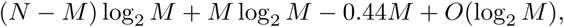

which is, ignoring the additive term *O*(log_2_ M), ≥ *N log_2_M — 0.44M*. Given the range of *M (i.e.,1 ≤ M ≤N)*, *N* log_2_ *M* always dominates the lower bound.

Now, we show that the space needed by Rainbowfish to store the variable-length labels assigned to color classes is equal to the lower bound. As explained in Theorem 1, the upper bound to store any label is log_2_*M* bits, and for *N* edges, it is given by *N* log_2_*M* bits. Rainbowfish also stores a boundary bit vector which has the same number of bits as the label bit vector. Therefore, the space required to store the label mappings is strictly ≤ 2*N* log_2_*M*. Note that the extra overhead to store the metadata to perform a select operation in constant time on the boundary bit vector is bounded by *o(N)*, where *N* is the numbers of bits in the bit vector [4].

However, Rainbowfish’s representation of color classes is entropy compressed (see Section 3.1) and the space required depends on the entropy of the color class distribution. For a highly skewed distribution, the entropy is low and the space required to store labels is much smaller than *N* log_2_*M* bits. On the other hand, when the distribution is near-uniform, i.e., the entropy is high, Rainbowfish makes all labels to be log_2_*M* bits and dispenses with the boundary bit vector. Therefore, the space required by Rainbowfish is always smaller than or equal to the lower bound.

### 3.4 Implementation

#### Storing bit vectors

In Rainbowfish, we use bit vector implementations from the SDSL library [3] to store the three bit vectors from Figure 1. We use the *rrr_vector* implementation from SDSL to store the equivalence class table and boundary bit vector, and the *bit_vector* implementation from SDSL to store the label bit vector.

The *rrr_vector* of SDSL is an implementation of RRR encoding [14]. RRR encoding is an entropy compressed encoding and also supports constant time rank and select operations on the compressed bit vector. The space reduction depends on the entropy of the bit vector. For high entropy bit vectors, the compression is not noticeable and in fact “negative” in some cases because of the extra metadata overhead to support rank and select operations.

The equivalence class table and boundary bit vector often have fairly low entropy, and can be compressed efficiently using RRR encoding. However, the label bit vector often has high entropy, and compressing it using RRR encoding is not effective. In our representation, the average order-0 entropy of the label bit vector for four different datasets is 0.94. This is a quite high, and hence we did not see any reduction in the space using RRR encoding. However, for the other two bit vectors, the order-0 entropy is lower (e.g., for boundary bit vector the average entropy over same four datasets is 0.56) and, in practice, we achieve a considerable space reduction using RRR encoding.

#### Construction

We use a 2-pass algorithm to construct the three bit vectors. In the first pass, we read the color matrix, compute the distinct color classes, and count the frequency of each class. Once we have the frequency information, we sort color classes in descending order based on their frequency. We then assign labels to color classes starting from zero. In the second pass, we read the uncompressed color matrix again, and add the label of each *k*-mer to the label bit vector. While building the label bit vector, we also build the boundary bit vector by storing a 1 at every index where a new label starts in the label bit vector. The labels are stored in the same order as the *k*-mers in the BOSS representation.

To reduce the space required for the labeling even further, we implemented our label encoding in the following way. Every time that the label size increases from *x* bits to *x* + 1 bits, we restart the counter of that label in label bit vector to 0. For example, we store 0 and 1 for labels 0 and 1 respectively, then we store 00, 01,10 and 11 for labels 2, 3, 4 and 5 respectively. For label value 6 we again restart the counter to 0 and store 000 to represent 6 in the label bit vector, etc. Later, when we want to retrieve the actual value of a label, we first recover the stored label *l’* from the label bit vector and then calculate the actual label *l* using the equation *l* = *l’ + 2^d^ –* 2 where *d* is length of label *l* in bits.

As explained in Section 3.2, the 2-pass algorithm minimizes the space used to represent color class labels by sorting the classes based on their frequencies and assigning labels to color classes to minimize the length of the resulting code path, similar to Huffman coding. However, one could also imagine assigning labels to color classes as we see them in the order *k*-mers appear in the BOSS representation. This way, we can construct all three tables in a single pass (i.e., a 1-pass algorithm).

However, as shown in Figure 2, this 1-pass algorithm can end up assigning long labels to frequent *k*-mers, and hence produce poor (i.e., large) encodings. However, the 2-pass algorithm always assigns labels according to the corresponding frequency distribution of the color classes. Sometimes, the 1-pass algorithm does well, but we chose to adopt the 2-pass algorithm in Rainbowfish.

**Figure 2.**
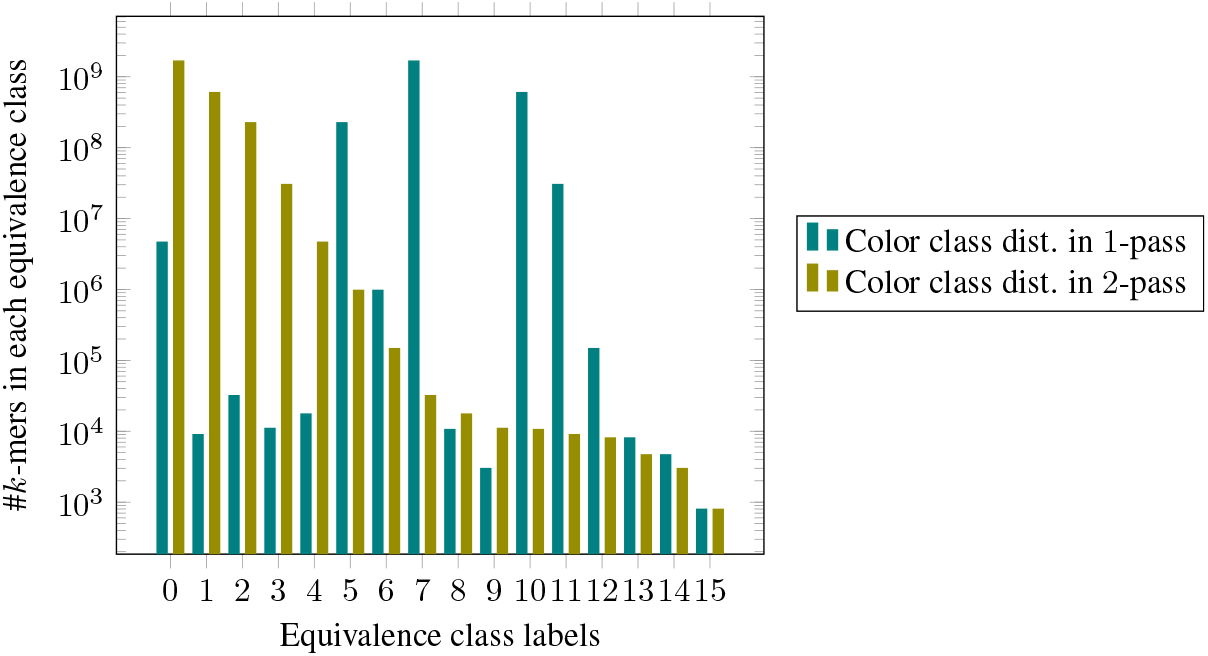
Distribution of *k*-mer frequencies across equivalence class labels in Rainbowfish after 1-*pass* and 2-*pass* algorithm on plant dataset Table 1. The 2-*pass* algorithm assigns the smallest label to color class with maximum number of *k*-mers. The distribution in 2-*pass* algorithm is monotonically decreasing.

## 4 Evaluation

In this section we evaluate Rainbowfish, and compare it to VARI [9], a state-of-the-art colored de Bruijn graph representation. We evaluate both systems in terms of space and running time. We address the following questions about the performance of Rainbowfish: How does Rainbowfish compare to VARI in terms of the space required to represent color information?; How does Rainbowfish compare to VARI in terms of the construction time?; How does Rainbowfish compare to VARI in terms of typical queries (e.g., in bubble calling)? We are particularly concerned with ensuring that Rainbowfish produces small encodings of the color information and remains practically efficient to query.

### 4.1 Experimental setup

To answer the above questions, we perform two different benchmarks. First, we evaluate the time taken to construct the color class representation. The construction time is the time taken to construct the color class representation from a list of color classes stored in the order of the edges in the de Bruijn graph (this is the same input used by VARI). During construction, we adopt a two-pass algorithm. In the first pass, we use a hash-table to determine the distinct color classes and the cardinality of each such class. Given this information, we know exactly the number of bits that will be required to store the label and boundary vectors. In the second pass, we fill in both the label and boundary vectors and then save all three structures to file. The construction time recorded here does not include (for either Rainbowfish or VARI) the time taken to build the de Bruijn graph and color list corresponding to edges in the de Bruijn graph (since this is the same for both methods).

We also report the space needed by both Rainbowfish and VARI to store the color class representation on disk. We do not include the space needed to represent the actual de Bruijn graph in our space comparisons because both Rainbowfish and VARI use BOSS to store the actual de Bruijn graph, and the BOSS representation itself tends to take less space than the color information.

Second, we evaluate the time taken to perform the bubble calling benchmark as described in [10], using both the VARI and Rainbowfish representations. Finding bubbles in a colored de Bruijn graph enables one to detect regions in the de Bruijn graph where different samples (i.e., colors) diverge from each other. As originally suggested by Iqbal et al. [6], such algorithms can form the basis for analyzing certain types of genetic variants in populations of genomes. We note that we adopt the exact bubble calling algorithm implemented in VARI, and the only variable being altered in our bubble-calling benchmark is the data structure being used to determine the set of colors present for each edge. Briefly, the bubble calling algorithm takes as input a pair *c_1;_ c_2_* of colors and traverses edges in the de Bruijn graph to find bubbles in which the edges in one sub-path are colored with *c*_1_ and the edges in the other sub-path are colored with *c*_2_ (see [10] for further details).

For all experiments in this paper, we consider the *k*-mer size to be 32. We carry out these benchmarks on a number of datasets as described in section 4.2. The time reported for construction and bubble calling are averaged over two runs, and the time is measured as the wall-clock time using the `/usr/bin/time` executable. All experiments were performed on an Intel(R) Xeon(R) CPU (E5-2699 v4@2.20GHz with 44 cores and 56MB L3 cache) with 512GB RAM and a 4TB TOSHIBA MG03ACA4 ATA HDD running ubuntu 16.10, and were carried out using a single thread.

### 4.2 Data

We run our benchmarks on the datasets mentioned in Table 1. The first three datasets, *E. coli*, Plant, and Beef safety are slight variants of those used for evaluation in VARI [10]. Each of these data sets exhibits different characteristics in terms of the number of *k*-mers, the number of input samples (i.e., colors) and the homogeneity of the underlying samples (i.e., how different are the de Bruijn graph for each of the individual samples). The first dataset consists of the assemblies of 5,598 different strains of *E. coli* obtained from GenBank [12]. Here, each “color” represents a specific *E. coli* assembly. Since these assemblies are from different strains of the same species, they exhibit a small degree of heterogeneity. In other words, a large fraction of the union de Bruijn graph is expected to occur in all samples.

**Table 1.** Different datasets used in our experiments. Number of edges include *k*-mers and dummy edges in the BOSS representation. *# of edges excluding dummies.

| Datasets | # of edges | # of colors (samples) | # of distinct color classes |
| --- | --- | --- | --- |
| *E. coli* 10 | 28,273,951 | 10 | 479 |
| *E. coli* 1000 | 157,737,064 | 1000 | 2,669,157 |
| *E. coli* 5598 | 435,705,390 | 5598 | 7,000,715 |
| Plant | 2,520,140,426 | 4 | 16 |
| Beef safety | 97,096,576,010* | 87 | 623,022,532 |
| Human transcriptome | 159,441,804* | 95,146 | 340,762 |

To evaluate the scalability of Rainbowfish when primarily changing the underlying number of input colors, we have evaluated three variants of the *E. coli* dataset. These consist of a dataset containing only 10 different strains, another containing 1,000 different strains and the final containing all 5,598 strains.

The second dataset (i.e., Plant) consists of the genome assemblies of four different plant species. Hence, this dataset contains only four colors, but has more than ≈ 2 billion distinct *k*-mers. The plant species considered are, A. *thaliana* ^1^ [16], Corn^2^ [15], Rice^3^ [17], and Tomato^4^ [2]. These genomes exhibit considerable diversity and heterogeneity. Given the diverse regions in the colored de Bruijn graph, this dataset is a good candidate for the bubble calling benchmark. Further, Muggli et al. [10] found that this was the only of the three original datasets on which they were able to construct the original `Cortex` representation of the colored de Bruijn graph. They validated that VARI produces the same bubble calls as `Cortex` here [10], and VARI and Rainbowfish, of course, produce the same bubble calls.

The third dataset, Beef safety, is considerably different from the prior data. Instead of the input samples consisting of assembled genomes, they consist of 87 metagenomic samples sequenced from cattle in the commercial process of beef production [11]. Hence, this dataset yields a considerably larger and more complex de Bruijn graph since it is built upon many un-assembled (and non-error-corrected) reads. Thus, the de Bruijn graph will encode portions of the relevant metagenomes as well as the effects of sequencing errors. This dataset also has many more *k*-mers than the others, ≈ 97 billion. It exhibits a large degree of heterogeneity and an intermediate number of input colors (87).

In addition to the three datasets used in the VARI paper, we also consider building the colored de Bruijn graph on the human transcriptome^5^ (Gencode v26 protein coding transcripts) [5]. Here, we consider each transcript as an individual sample (i.e., a distinct input color). This data consists of ≈ 95, 000 colors, but only ≈ 159 million *k*-mers. Hence, this dataset will give an idea about how the representations will perform when the number of colors becomes very large (though the number of distinct color classes remains orders of magnitude smaller than the number of *k*-mers). Further, we note that this dataset highlights some of the similarities between the color class encoding adopted by Rainbowfish and the k-mer-based equivalence class decomposition adopted by certain transcript quantification methods (e.g. [13]).

### 4.3 Performance

Table 2 shows the time taken by Rainbowfish and VARI to construct the color class representation for different datasets. Rainbowfish uses a 2-pass algorithm to construct the color class representation, and hence the construction time is dominated by the steps to read the color list file twice. For small datasets like *E. coli* 10 and *E. coli* 1, 000, the input file size is small and does not affect the overall construction time compared to VARI. However, for large datasets like Plant and Beef safety, the time to read the color file twice dominates the construction time and Rainbowfish is 1.9 ⨯—3⨯ slower. We note that this time can be considerably reduced by avoiding the uncompressed color matrix representation currently used upstream of Rainbowfish and VARI, and integrating determination and encoding of the color classes into the de Bruijn graph construction directly. However, this is outside the scope of the current paper.

**Table 2.** Construction and bubble calling time for Rainbowfish and VARI for different datasets.

| Datasets | Construction Time (secs) |  | Bubble Calling Time (secs) |  |
| --- | --- | --- | --- | --- |
|  | VARI | Rainbowfish | VARI | Rainbowfish |
| *E. coli* 10 | 44 | 31 | 344 | 366 |
| *E. coli* 1000 | 340 | 270 | 2,610 | 2,356 |
| *E. coli* 5598 | 3,141 | 4,021 | 8,796 | 8,201 |
| Plant | 108 | 339 | 47,040 | 48,537 |
| Beef safety | 15,378 | 30,478 | NA | NA |
| Human transcriptome | 13,961 | 30,804 | NA | NA |

#### Space

Table 3 shows the space usage of Rainbowfish and VARI for the different datasets we consider. Among these data, there are a range of characteristics in terms of the number of *k*-mers, the number of colors, and the complexity and heterogeneity of the de Bruijn graph. We find that, for all datasets, Rainbowfish requires less space to store the color information than VARI. The magnitude of the improvement depends on the makeup of the data, but is as large as ~ 20⨯.

**Table 3.** The space required by Rainbowfish and VARI to store the color class representation for different datasets. The first column shows space required for the uncompressed color matrix *(N ⨯ C bits)*. All space is reported in MB.

| Datasets | uncompressed color matrix | VARI | Rainbowfish |
| --- | --- | --- | --- |
| *E. coli* 10 | 34 | 58 | 20 |
| *E. coli* 1000 | 18,804 | 8,848 | 475 |
| *E. coli* 5598 | 290,761 | 58,718 | 2,938 |
| Plant | 1,202 | 1,603 | 497 |
| Beef safety | 1,007,009 | 210,998 | 144,564 |
| Human transcriptome | 1,808,435 | 841 | 817 |

In particular, Rainbowfish’s space usage is particularly impressive for datasets with a large number of input colors but a relatively small number of distinct *k*-mers. In this case, we usually find that the number of distinct color classes is very small compared to the universe of possibilities, and so each label can be encoded in much fewer than *C* bits. However, the space VARI consumes depends greatly on the sparsity of the color matrix. The color matrix itself grows rapidly as the number of *k*-mers and colors increases, but VARI’s compression mechanism (Elias-Fano encoding) is very effective if the color matrix is sparse (e.g., each *k*-mer is labeled with only a small subset of colors). This is exactly the case for the Human transcriptome, where the color matrix has an entropy of ~ 0.0004 (compared to *E. coli* 5,598 and *E. coli* 1,000 with entropies of ~ 0.16 and ~ 0.34 respectively). Thus, in the *E. coli* dataset, VARI can save space up to a factor of ~ 5 compared with the uncompressed representation, while in the Human transcriptome it can save a factor of ~ 2,150 because of the low entropy of the color matrix. Rainbowfish does well in all experiments, even when the number of input colors is small (e.g., in the Plant dataset). Rainbowfish achieves the most impressive compression when the color class distribution has low entropy and the number of color classes is small relative to the upper bound. In such cases, the entropy compressed representation of Rainbowfish is able to represent a large fraction of all labels using a very small number of bits.

#### Bubble calling

Table 2 shows the time taken by Rainbowfish and VARI to perform the bubble calling benchmark on different datasets. We run the bubble calling benchmark on the *E. coli* and Plant datasets (as in the VARI paper). We note that the current bubble calling algorithm is too slow to run on the Beef safety data set (the time in [10] was estimated at > 3, 000 hours). It is possible, however, that optimizations to the underlying algorithm might lift this restriction. We also did not perform bubble calling on the human transcriptome dataset as here, we were unable, given the resources on our server, to even run the de Bruijn graph construction to completion. Specifically, due to the large amount of external memory that VARI uses to build the uncompressed color matrix and the de Bruijn graph on these larger (either in terms of the number of *k*-mers, the number of colors, or both) datasets (on order of Terabytes), we exhausted the available disk space. For these datasets, to approximate the relevant sizes and construction times, we produced a uncompressed color matrix that lists the colors for each *k*-mer and its reverse complement, and we use this to build both the VARI and Rainbowfish color representations. While very similar to the full color matrix that VARI would produce, this file is slightly different in that it does not include entries for dummy edges (a detail of the BOSS representation), and the order of the color matrix rows can be different from what will appear in the BOSS representation. However, we still believe these numbers, provided in Table 1, give a reasonable approximation of how the respective methods would perform were we able to construct the de Bruijn graph completely.

For bubble calling, both representations require a very similar amount of time. This is likely due, in part, to the fact that navigating the BOSS representation of the de Bruijn graph may be the performance bottleneck in the bubble calling algorithm. Thus, both VARI and Rainbowfish provide sufficiently fast access to the color sets for each edge that they do not represent bottlenecks in this regard.

## 5 Conclusion and Future Work

In this paper, we propose an entropy-compressed, succinct data structure to store the color information of a colored de Bruijn graph. To represent the topology of the de Bruijn graph itself, we adopt the BOSS [1] representation. However, we note that, for our representation of the color sets, we only require that the underlying de Bruijn graph representation is able to associate a unique rank between 0 and *N* — 1 with each edge. Hence, it is possible to use the Rainbowfish representation with other representations of the de Bruijn graph topology (e.g., those based on minimal perfect hashing).

We demonstrate that the inherent skewness in the distribution of color classes can be exploited to reduce the size of the color information. This allows Rainbowfish to represent the colored de Bruijn graph, even for large datasets with many colors, in a reasonably small space. In fact, for representing the color information itself, we show that Rainbowfish is succinct, and hence requires only *Z* + *o(z)* bits where *Z* is the number of bits required by an information theoretic optimal representation.

While we have described here a system for efficiently representing the color information in a colored de Bruijn graph, our encoding scheme can be generalized to store any type of attribute attached to the edges. For example, the one could use the same (or a related) scheme to encode information like the *k*-mer count or set of positions associated with a given edge. Moreover, it will be interesting to explore how multiple attributes could be efficiently stored simultaneously, and how potential correlations between these attributes might be exploited.

Finally, in our current implementation, the input to the system is a color matrix file generated by VARI. This implementation requires first building the uncompressed color matrix, and then permuting the rows of this matrix along with the edges of the de Bruijn graph during the BOSS construction procedure.

This process can require a large amount of space, as the uncompressed color matrix can become extremely large (on the order of Terabytes for some of the datasets we considered here). Consequently, in most cases, the construction algorithm must resort to making extensive use of external memory (i.e., disk), which increases building time and consumes a lot of disk space. However, by building the Rainbowfish representation of the color sets prior to BOSS construction, and simply associating the relevant labels, rather than uncompressed color vectors, with each edge, it is likely possible to vastly reduce the time and spaced required to construct the colored de Bruijn graph. Thus, in the future, we are interested in both incorporating the Rainbowfish representation more tightly inside the existing VARI codebase, as well as pairing the Rainbowfish representation with other compatible representations of the de Bruijn graph topology.

## Acknowledgments

We gratefully acknowledge support from NSF grant BBSRC-NSF/BIO-1564917. We also thank Michael Bender and Robert Johnson for fruitful conversations and important insight when performing this research.

## Footnotes

1 ftp://ftp.ensemblgenomes.org/pub/plants/release-34/fasta/arabidopsis_thaliana/dna/Arabidopsis_thaliana.TAIR10.dna.toplevel.fa.gz

2 ftp://ftp.ncbi.nlm.nih.gov/genomes/all/GCF/000/005/005/GCF_000005005.1_B73_RefGen_v3/GCF_000005005.1_B73_RefGen_v3_genomic.fna.gz

3 http://rice.plantbiology.msu.edu/pub/data/Eukaryotic_Projects/o_sativa/annotation_dbs/pseudomolecules/version_7.0/all.dir/all.con

4 ftp://ftp.solgenomics.net/tomato_genome/assembly/build_2.50/SL2.50ch00.fa.

5 ftp://ftp.sanger.ac.uk/pub/gencode/Gencode_human/release_26/gencode.v26.pc_transcripts.fa.gz

## References

1. Alexander Bowe, Taku Onodera, Kunihiko Sadakane, and Tetsuo Shibuya. Succinct de Bruijn graphs. In Proceedings of the International Workshop on Algorithms in Bioinformatics, pages225–235. Springer, 2012.

2. Mathilde Causse, Nelly Desplat, Laura Pascual, Marie-Christine Le Paslier, Christopher Sauvage, Guillaume Bauchet, Aurelie Berard, Remi Bounon, Maria Tchoumakov, Dominique Brunel, et al. Whole genome resequencing in tomato reveals variation associated with introgression and breeding events. BMC genomics, 14(1):791, 2013.

3. Simon Gog. Succinct data structure library. https://github.com/simongog/sdsl-lite, 2017. [online; accessed 01-Feb-2017].

4. Rodrigo Gonzalez, Szymon Grabowski, Veli Mäkinen, and Gonzalo Navarro. Practical implementation of rank and select queries. In Poster Proceedings Volume of 4th Workshop on Efficient and Experimental Algorithms (WEA), pages 27–38, 2005.

5. J. Harrow,A. Frankish, J. M. Frankish, J. M. Gonzalez, E. Tapanari, M. Diekhans, F. Kokocinski, B. L. Kokocinski, B. L. Aken, D. Barrell, A. Zadissa, S. Searle, I. Barnes, A. Bignell, V. Boychenko, T. Hunt, M. Kay, G. Mukherjee, J. Rajan, G. Despacio-Reyes, G. Saunders, C. Steward, R. Harte, M. Lin, C. Howald, A. Tanzer, T. Derrien, J. Chrast, N. Walters, S. Balasubramanian, B. Pei, M. Tress, J. M. Tress, J. M. Rodriguez, I. Ezkurdia, J. van Baren, M. Brent, D. Haussler, M. Kellis, A. Valencia, A. Reymond, M. Gerstein, R. Guigo, and T. J. Hubbard. GENCODE: The reference human genome annotation for the ENCODE project. Genome Research, 22(9):1760–1774, sep 2012. URL:https://doi.org/10.1101%2Fgr.135350.111,doi:10.1101/gr.135350.111.

6. Zamin Iqbal, Mario Caccamo, Isaac Turner, Paul Flicek, and Gil McVean. De novo assembly and genotyping of variants using colored de Bruijn graphs. Nature genetics, 44(2):226–232, 2012.

7. Guy Jacobson.Space-efficient static trees and graphs. In Foundations of Computer Science, 1989., 30th Annual Symposium on, pages 549–554. IEEE, 1989.

8. Guy Joseph Jacobson. Succinct Static Data Structures. PhD thesis, Carnegie Mellon University, Pittsburgh, PA, USA, 1988. AAI8918056.

9. Muggli Martin D. Vari. https://github.com/cosmo-team/cosmo/tree/VARI, February 2017. Viewed Feb 3, 2017.

10. Martin D. Muggli, Alexander Bowe, Noelle R. Noyes, Paul Morley, Keith Belk, Robert Raymond, Travis Gagie, Simon J. Puglisi, and Christina Boucher. Succinct Colored de Bruijn Graphs. Bioinformatics, 2017.

11. Noelle R Noyes, Xiang Yang, Lyndsey M Linke, Roberta J Magnuson, Adam Dettenwanger, Shaun Cook, Ifigenia Geornaras, Dale E Woerner, Sheryl P Gow, Tim A McAllister, et al. Resistome diversity in cattle and the environment decreases during beef production. ELife, 5:e13195, 2016.

12. Nuala A O’Leary, Mathew W Wright, J Rodney Brister, Stacy Ciufo, Diana Haddad, Rich McVeigh, Bhanu Rajput, Barbara Robbertse, Brian Smith-White, Danso Ako-Adjei, et al. Reference sequence (refseq) database at ncbi: current status, taxonomic expansion, and functional annotation. Nucleic acids research, page gkv1189, 2015.

13. Rob Patro, Stephen M Mount, and Carl Kingsford. Sailfish enables alignment-free isoform quantification from RNA-seq reads using lightweight algorithms. Nature biotechnology, 32(5):462–464, 2014.

14. Rajeev Raman, Venkatesh Raman, and S Srinivasa Rao. Succinct indexable dictionaries with applications to encoding k-ary trees and multisets. In Proceedings of the thirteenth annual ACM-SIAM symposium on Discrete algorithms, pages 233–242. Society for Industrial and Applied Mathematics, 2002.

15. Patrick S Schnable, Doreen Ware, Robert S Fulton, Joshua C Stein, Fusheng Wei, Shiran Pasternak, Chengzhi Liang, Jianwei Zhang, Lucinda Fulton, Tina A Graves, et al. The b73 maize genome: complexity, diversity, and dynamics. science, 326(5956):1112–1115, 2009.

16. David Swarbreck, Christopher Wilks, Philippe Lamesch, Tanya Z Berardini, Margarita Garcia-Hernandez, Hartmut Foerster, Donghui Li, Tom Meyer, Robert Muller, Larry Ploetz, et al. The arabidopsis information resource (tair): gene structure and function annotation. Nucleic acids research, 36(suppl 1):D1009 –D1014, 2008.

17. Tsuyoshi Tanaka, Baltazar A Antonio, Shoshi Kikuchi, Takashi Matsumoto, Yoshiaki Nagamura, Hisataka Numa, Hiroaki Sakai, Jianzhong Wu, Takeshi Itoh, Takuji Sasaki, et al. The rice annotation project database (rap-db): 2008 update. Nucleic Acids Research, 36(Supp 1):D1028–D1033, 2008.

